# DOT1L methyltransferase regulates the calcium influx in erythroid progenitor cells in response to erythropoietin

**DOI:** 10.1101/2020.10.04.325746

**Authors:** Yi Feng, Shaon Borosha, Anamika Ratri, Sami M. Housami, V. Praveen Chakravarthi, Huizhen Wang, Jay L. Vivian, Timothy A Fields, William Kinsey, MA Karim Rumi, Patrick E. Fields

## Abstract

Erythropoietin (EPO) signaling plays a vital role in erythropoiesis by regulating proliferation and lineage-specific differentiation of hematopoietic progenitor cells. An important downstream response of EPO signaling is calcium influx, which is regulated by transient receptor potential channel (TRPC) proteins, particularly TRPC2 and TRPC6. While EPO induces Ca^2+^influx through TRPC2, TRPC6 inhibits the function of TRPC2. Thus, interactions between TRPC2 and TRPC6 regulate the rate of Ca^2+^influx in EPO-induced erythropoiesis. In this study, we observed that the expression of TRPC6 in c-KIT positive erythroid progenitor cells is regulated by DOT1L. DOT1L is a methyltransferase that plays an important role in many biological processes during embryonic development, including early erythropoiesis. We previously reported that *Dot1L* knockout (*Dot1L-KO*) hematopoietic progenitors in the yolk sac failed to develop properly, which resulted in lethal anemia. In this study, we have detected a marked downregulation of *Trpc6* gene expression in *Dot1L-KO* progenitor cells in the yolk sac compared to wildtype. However, the expression of *Trpc2*, the positive regulator of Ca^2+^influx, remained unchanged. The promoter and the proximal region of the *Trpc6* gene loci exhibited an enrichment of H3K79 methylation, which is mediated solely by DOT1L. As the loss of DOT1L affects the expression of TRPC6, which inhibits Ca^2+^influx by TRPC2, *Dot1L-KO* progenitor cells in the yolk sac exhibit accelerated and sustained high levels of Ca^2+^influx. Such heightened Ca^2+^ levels might have detrimental effects on the development of hematopoietic progenitor cells in response to erythropoietin.

## 1. INTRODUCTION

Regulation of erythropoiesis is a highly orchestrated process involving epigenetic and transcriptional regulation in response to a number of cytokines and growth factors (Tsiftsoglou et al., 2009). Amongst all external factors, erythropoietin (EPO) is considered the primary regulator of erythropoiesis (Ingley et al., 2004). When EPO binds to EPO receptors (EPOR), several signaling pathways are initiated, particularly the PLCγ signaling pathway (Tsiftsoglou et al., 2009). Activation of PLCγ generates the second messenger inositol-1,4,5-trisphosphate (IP3). Studies have shown that a conformational change of the IP3 receptor upon IP3 binding facilitates its association with the transient receptor potential channel (TRPC) proteins, which are voltage-independent Ca^2+^channels (Fisher, 2003; Miller et al., 1994; Tong et al., 2004; Tsiftsoglou et al., 2009). TRPC proteins mediate Ca^2+^influx into erythroid progenitors (Tong et al., 2004; Tong et al., 2008). It has been demonstrated that regulation of intracellular Ca^2+^by EPO plays a critical role in the survival, proliferation, and differentiation of erythroid progenitors (Hensold et al., 1991; Miller et al., 1989). Among the six members of the mouse TRPC family, TRPC2 and TRPC6 mRNAs and proteins are expressed in erythropoietic cell lines (Tong et al., 2004; Tong et al., 2008). EPO stimulation of erythroid cells induces the Ca^2+^influx through TRPC2, while TRPC6 inhibits the function of TRPC2. Therefore, the interaction of TRPC2 and TRPC6 plays an important role in hematopoietic cells to regulate Ca^2+^influx in response to EPO stimulation.

Disruptor of telomeric silencing 1-like (DOT1L) plays a crucial role in many embryonic developmental processes (Feng et al., 2002; Feng et al., 2010; Malcom et al., 2020). DOTL1L is the only known methyltransferase in eukaryotic cells that is responsible for the mono, di- and tri-methyl marks on lysine 79 of histone H3 (H3K79) (Feng et al., 2002), and these histone modifications are strongly associated with actively transcribed chromatin regions (Steger et al., 2008). We previously reported that DOT1L deficiency in mutant (*Dot1L-KO*) mice resulted in lethal anemia during mid-gestation (Feng et al., 2010). *Dot1L-KO* erythroid progenitors failed to develop normally, showing cell cycle arrest and increased apoptosis (Feng et al., 2010). However, the molecular mechanisms underlying DOTL1L regulation of early erythropoiesis remain unclear. In this study, we observed that *Trpc6* is a direct target of DOTL1L in hematopoietic progenitor cells (HPCs) isolated from yolk sacs. We detected an enrichment of H3K79 methylation within the *Trpc6* gene loci. Moreover, loss of *Trpc6* expression in *Dot1L-KO* HPCs showed an accelerated and sustained high levels of Ca^2+^influx.

## 2. MATERIALS AND METHODS

### 2.1. Mouse lines and isolation of cells from yolk sac

The *Dot1L-*KO mouse line was generated in our previous study (Feng et al., 2010) and maintained by continuous backcrossing into C57BL/6 strains. For all the assays in this study, HPCs were derived from yolk sacs on embryonic day 10.5 (E10.5). Single-cell suspension was obtained as described before (Feng et al., 2010). Briefly, yolk sacs were incubated in 0.1% collagenase at 37°C for 30 minutes. Then the yolk sacs were aspirated through 25-G and 27-G needles sequentially and filtered through a 70 µM strainer. Genotyping was performed by PCR using DNA extracted from corresponding embryo tissues (Feng et al., 2010). All animal experiments were performed in accordance with the protocols approved by the University of Kansas Medical Center (KUMC) Animal Care and Use Committee.

### 2.2. Assessment of cell proliferation, cell cycle analyses and apoptosis assays

Single-cell suspensions from E10.5 yolk sacs were cultured in M3334 medium and cell proliferation was assessed by standard cell-counting by hemocytometer on every alternate days of culture (Feng et al., 2010). Prior to cell cycle studies, E10.5 HPCs were cultured in M3334 medium for 4 days to generate erythroid colonies. The cells were separated by gentle pipetting, fixed by adding cold 70% ethanol slowly to the cell suspensions, and treated with RNase. Cells were then stained with propidium iodide and analyzed by flow cytometry for cell cycle progression (Feng et al., 2010; Malcom et al., 2020). Separately, fixed erythroid cells were labeled with Annexin V and then analyzed by flow cytometry for signs of apoptosis as described previously (Malcom et al, 2020). Flow cytometry was performed using FACSCalibur (BD Biosciences, San Jose, CA) and analyses of cytometric data was carried out using CellQuest Pro software (BD Biosciences) ((Feng et al., 2010; Malcom et al., 2020).

### 2.3. Isolation of c-KIT Cell positive cells yolk sac

Single-cell suspensions were prepared from E10.5 yolk sacs as described above and c-KIT positive cells were separated as done previously (Feng et al, 2010). Briefly, HPCs were incubated with an anti-c-KIT antibody conjugated to phycoerythrin cyanine (eBioscience Inc, 25-4317-82) at 4°C for 30 minutes. Then the c-KIT positive cells were isolated by cell sorting using a BD FACSAria cell sorter (BD Bioscience) at the KUMC flow cytometry core.

### 2.4. RNA extraction, cDNA preparation and RT-qPCR

Total RNA was extracted using TRIzol reagent (ThermoFisher Scientific) following the manufacturer’s instructions. cDNAs were prepared from 1µg of total RNA collected from each sample using SuperScript VILO cDNA Synthesis Kit (ThermoFisher Scientific). Quantitative real-time PCR (qPCR) was performed using Power SYBR Green Master Mix (ThermoFisher Scientific) and ran on a 7500 real-time PCR system (Applied Biosystems). qPCR results were normalized to Rn18S expression and calculated by the comparative ΔΔCT method (Chakravarthi et al., 2020; Khristi et al., 2018; Khristi et al., 2019). A list of qPCR primer sequences used in this study is shown in **Table 1** (http://pga.mgh.harvard.edu/primerbank/).

**Table 1:**
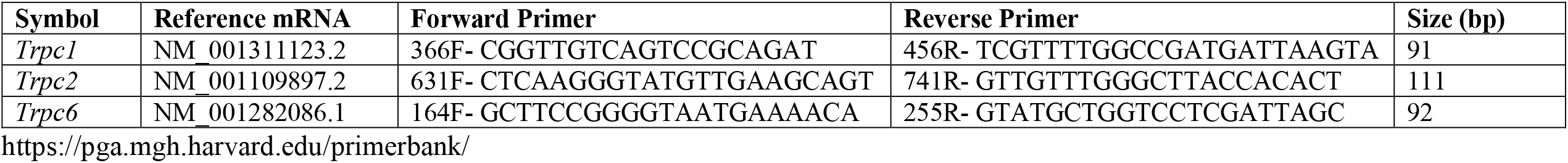
Primers used in qRT-PCR studies

### 2.5. Western blot analyses

Western blot analyses of TRPC6 were performed following standard protocols. Cells from the whole yolk sac, bone marrow, or liver were lysed in Nonidet-P40 (NP-40) buffer (150 mM sodium chloride, 1.0% NP-40, 50 mM Tris, pH 8.0) on ice for 30 minutes. Cell extracts were centrifuged at 12000g at 4°C for 20 minutes. Supernatants were collected and protein concentrations were measured using Bradford assay (BioRad). Approximately 30µg of each protein sample was mixed with NuPAGE SDS 4X sample buffer (ThermoFisher Scientific) and were electrophoresed on a 10% precast gel in SDS running buffer (250 mM Tris base, 190 mM glycine, 0.1% SDS, pH8.3) (BioRad). Proteins were transferred to PVDF membranes and the membranes were blocked in 5% bovine serum albumin (BSA) overnight at 4°C. Blocked membranes were incubated with 1:300 TRPC6 primary antibody (Abcam, mAbcam63038) for 4 hours at 4°C with shaking. Membranes were washed 5 times with TBST buffer at 4°C with shaking and incubated with 1:5000 secondary antibody (Goat anti-mouse IRDye CW800) for 45 minutes at room temperature. Then the membranes were washed another 5 times with TBST, and images were captured with an Odyssey CLx infrared image system (LI-COR Biosciences). The same membrane was stripped and incubated with an antibody against ACTB (Millipore Sigma) as loading control of protein samples.

### 2.6. Extraction of bone marrow and liver cells

Bone marrow cells that express high levels of *Trpc6* (Ichikawa and Inoue, 2014) were isolated from adult male mouse femur as described previously (Ichikawa and Inoue, 2014). Briefly, skin and muscles from the pelvic and femoral bones were removed. The bones were cleaned, and the ends were cut off at each end. Bone marrows were expelled from the ends of the bone passing cold sterile RNAse free PBS with a 5ml syringe fitted with a 25G needle. Then the cell aggregates were dispersed, and marrow cells were separated by repeated aspiration with a 5ml syringe attached with 27G needle and passing through a 70µM strainer. For liver cell isolation, mouse liver tissues were dissected and minced into small pieces in presence cold sterile RNAse free PBS. The tissue pieces were ground between glass slides and passed through 70µM strainer. Strained bone marrow or liver cells were washed with cold, sterile RNAse free PBS used for RNA or protein extraction or fixed in formaldehyde for ChIP assays.

### 2.7. ChIP assay for H3K79 Di- and Tri-methylation in Trpc6 locus

Chromatin immunoprecipitation (ChIP) was carried out following standard protocols (Chakravarthi et al., 2018; Chuong et al., 2013). Briefly, for each assay, 10^7^ cells were resuspended in 10ml IMDM medium containing 5% FBS and were fixed in 1% formaldehyde for 10min. Glycine was added to a final concentration of 125mM and incubated at room temperature with rotation for 10min to neutralize formaldehyde. The cells were washed twice with cold PBS, lysed in 500µl SDS lysis buffer with protease inhibitor cocktail (Millipore Sigma), and incubated on ice for 10min. The nuclear pellets were washed with a cold lysis buffer and sonicated using a probe sonicator (Branson Sonifier 250) 4-6 times, 15sec each time, to reduce the chromatin length to between 200 and 1000bp. Sonicated chromatin solutions were centrifuged at 10,000g at 4°C for 10min. Clear supernatants were collected into 1.5ml tubes and ∼500µl chromatin solution was mixed with 75µl Protein A agarose/Salmon Sperm DNA and incubated at 4°C with rotation overnight. The solution was centrifuged 3000g for 5min. The supernatant was transferred to a new tube and diluted 10-fold in ChIP Dilution Buffer. 2µg of H3K79me2, or H3K79me3 antibody (Abcam Inc) was added to 1ml of the diluted chromatin solution each and another 1ml was used as ‘input’. 2µg of normal rabbit IgG (BioSource International) was used as negative control. Chromatin solutions were incubated at 4°C with rotation for 4h in presence of specific antibodies. Then 65ul Protein A agarose/Salmon Sperm DNA was added to each tube and incubated at 4°C with rotation for another 2h. The solutions were centrifuged at 3000g for 5min at 4°C and supernatants were discarded. The Protein A agarose beads were washed for 5min on a rotating platform sequentially with 1ml of each of the Low Salt Wash Buffer, High Salt Wash Buffer, LiCl Wash Buffer, and 1XTE. Then the chromatins were eluted with 250µl elution buffers twice and elutes were combined. 20µl 5M NaCl and 2 µl RNase were added to each eluted chromatin samples and incubated at 37°C for 30min to remove RNAs and then again at 65°C for 4hto decrosslink the ChIPed chromatin materials. The chromatin solutions were mixed with 10µl 0.5M EDTA, 20µl 1M Tris-HCl, pH6.5, and 2µl 10mg/ml Proteinase K and incubated at 45°C for 1 hour. Finally, the chromatin DNA fragments were recovered by phenol/chloroform extraction and ethanol precipitation. Real-time PCR was performed to quantify precipitated DNA with the use of the 7500 real-time PCR system. ChIP-qPCR primers used for analyzing the *Trpc6* loci are listed in **Table 2**.

**Table 2:**
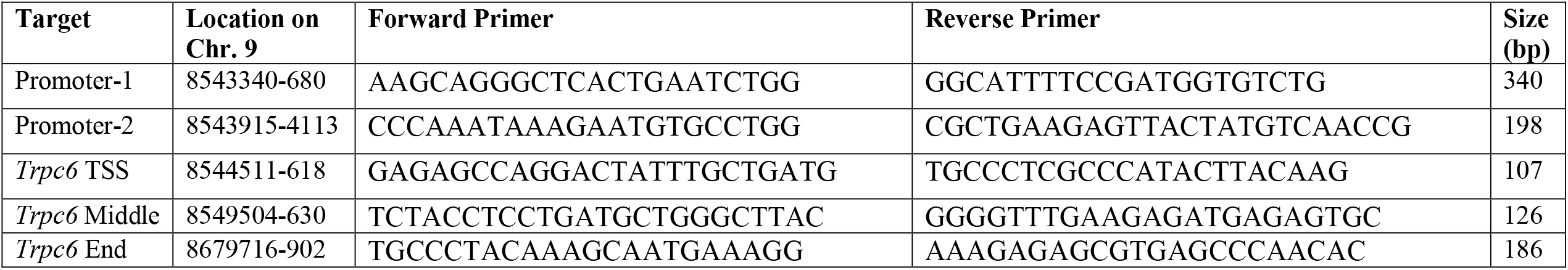
Primers for ChIP-qPCR for the promoter, TSS, and intragenic sites within the mouse *Trpc6* gene loci

### 2.8. Measuring calcium influx of yolk sac cells in response to EPO

A fluorescence microscopy-coupled digital imaging system was used to measure Ca2+ concentration change of HPCs when treated with EPO. Fura2 was used as the detection fluorophore. After the whole yolk sacs were dissociated into single cells, the cells were washed 2-3 times with PBS to completely remove fetal bovine serum (FBS). HPCs were resuspended in 40ul RPMI 1640 (Sigma) without FBS. Cells were loaded into the center of culture dish coated with poly-lysine and fixed for 10min at room temperature. The medium was then removed, and cells were stained with 1µM Fura2 and Pluronic F127 (Invitrogen) (1:1 in volume) at 37°C for 30min. Staining solution was removed, and cells were covered with 150µl RPMI containing 10% FBS. Mouse EPO (R&D systems) was added to a final concentration of 10 units/ml. Then the sample was analyzed immediately with the confocal fluorescence microscopy on a Nikon TE2000U microscope. Fura2-loaded cells were visualized with digital video imaging, and fluorescence was quantitated using the intensity ratio of the emission (510nm), which was measured following excitation at 340nm divided by the emission following excitation at 380nm. Each sample was measured over 20min. After that, the data was calculated and analyzed by Metamorph 6.1 (Universal Imaging Corp., Downingtown, PA).

### 2.9. Statistical analyses

Each experimental group consisted of a minimum of 6 samples. The experimental results are presented as the mean ± standard error (SE). The results were analyzed for one-way ANOVA, and the significance of mean differences was determined by Duncan’s post hoc test, with p < 0.05. All the statistical calculations were done using SPSS 22 (IBM, Armonk, NY).

## 3. RESULTS

### 3.1. Dot1L-KO erythroblasts displayed decreased proliferation, cell cycle arrest and increased apoptosis

Cells isolated from the E10.5 *Dot1L-KO* and wildtype yolk sacs were cultured in expansion medium and differentiated into ESREs (**Fig. 1A**). *Dot1L-*KO erythroblasts had severely blunted proliferation (**Fig. 1C)**, which resulted in a reduced number of growing cells compared to wildtype cells (**Fig. 1B**). We also detected that a large number of erythroid progenitors derived from *Dot1L-* KO yolk sacs displayed G0/G1 cell cycle arrest (**Fig. 1D-F**). An increased number of *Dot1L-*KO erythroid progenitors were also found to be apoptotic (**Fig. 1G**).

**Fig. 1.**
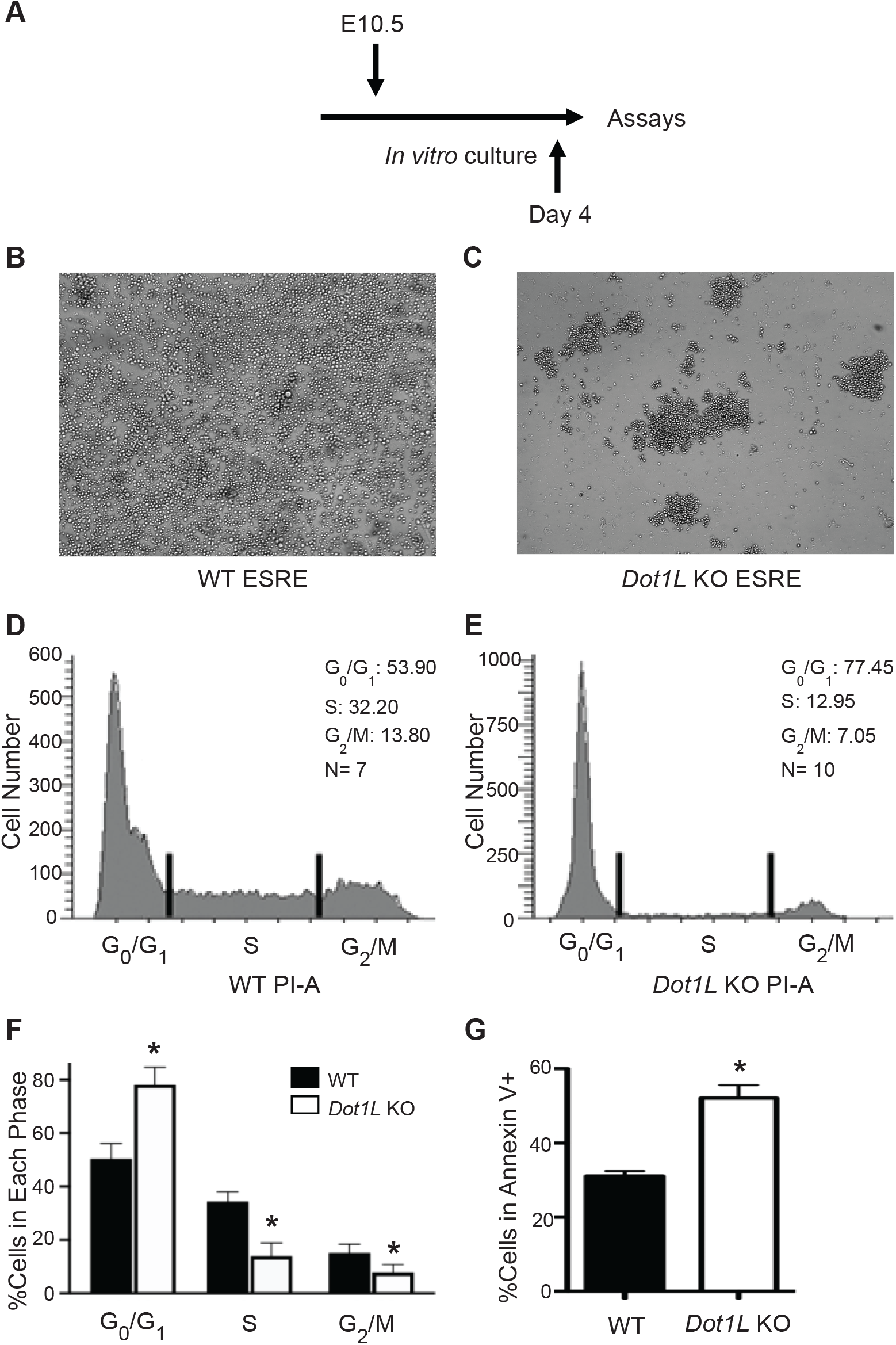
*Dot1L*-KO progenitors fail to proliferate in response to EPO. (**A**) Schematic diagram showing the timeline of cell cycle progression and apoptosis analysis. Cells were isolated from E10.5 KO and WT yolk sacs, cultured in expansion medium, and differentiated into ESREs. On day 4 of culture, cells were separated, fixed, and either stained with propidium iodide for cell cycle analyses or labeled with Annexin V for signs of apoptosis. Compared to WT ESREs (**B**), *Dot1L-* KO erythroblasts had severely reduced number of growing cells (**C**). A large number of erythroid progenitors from KO yolk sacs displayed G_0_/G_1_ arrest (**D-F**). In addition, an increased number of *Dot1L-*KO erythroid progenitors were found to be apoptotic (**G**). Data are expressed as mean±SE, n ≥ 6, * P< 0.05.

### 3.2. TRPC6 expression is downregulated in Dot1L-KO hematopoietic progenitor cells

We analyzed gene expression levels of c-KIT positive wildtype and *Dot1L-*KO hematopoietic progenitor cells isolated from E10.5 yolk sac. We observed that among three *Trpc* members, *Trpc1, Trpc2*, and *Trp6*, only *Trpc6* level was significantly reduced in *Dot1L-*KO progenitor cells (**Fig. 2A**). Other *Trpc* members remained relatively unchanged (**Fig. 2A**). We also detected a marked reduction in TRPC6 protein level in *Dot1L*-KO yolk sacs (**Fig. 2B** and **C**). These data indicate that DOT1L deficiency leads to reduced TRPC6 level in hematopoietic progenitor cells. Since TRPC2 level is not affected, it results in an increased TRPC2/TRPC6 ratio from 0.95 to 4.40.

**Fig. 2.**
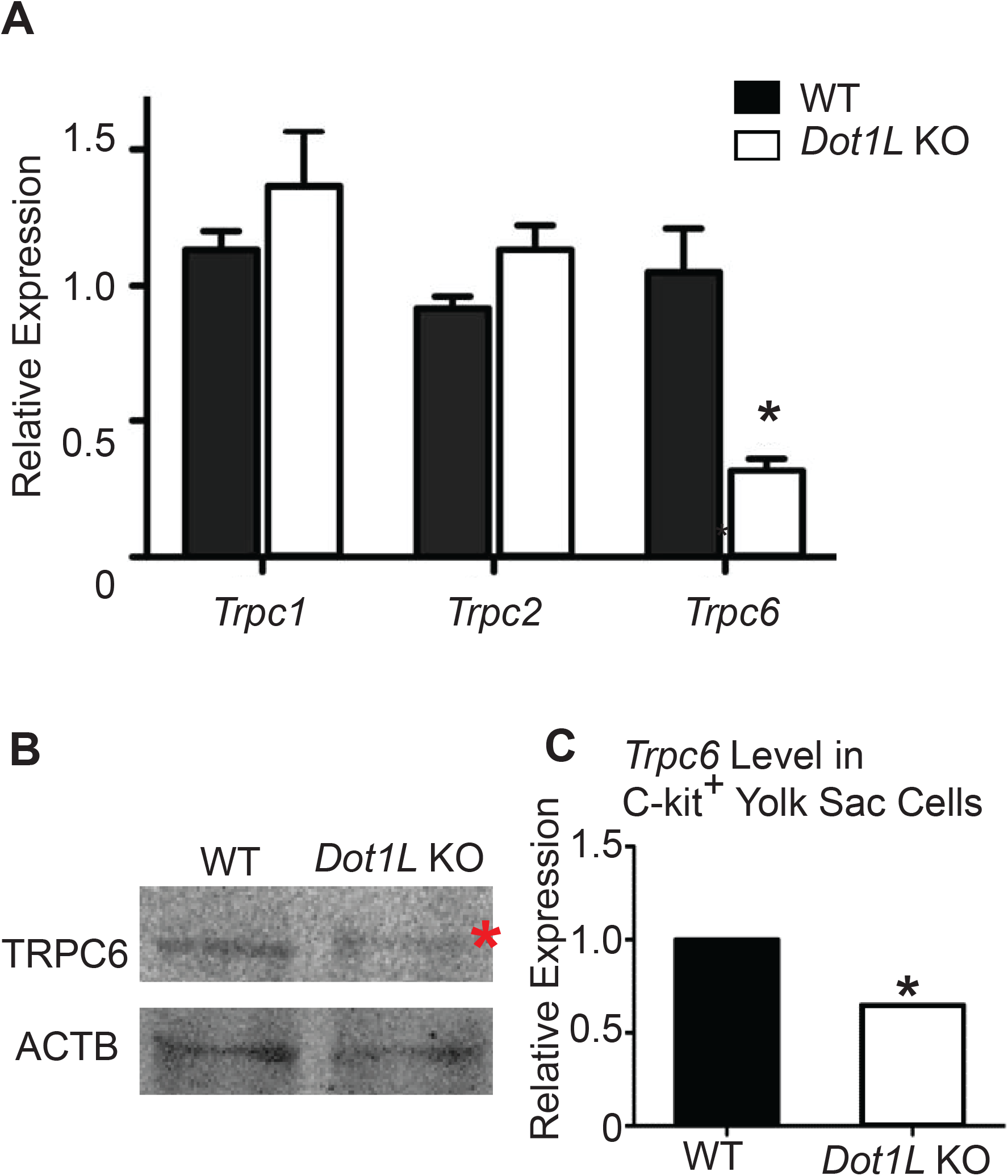
Expression of Trpc6 is markedly reduced in c-KIT positive *Dot1L*-KO hematopoietic progenitors. (**A**) Among three genes in the *Trpc* family, *Trpc1, Trpc2*, and *Trp6*, only *Trpc6* level was dramatically reduced in *Dot1L-*KO compared to WT, while the levels of the other members stayed relatively unchanged. TRPC6 protein level in c-KIT positive progenitor cells from KO yolk sacs was also significantly reduced (**B** and **C**). Data are expressed as mean±SE, n ≥ 6, * P< 0.05.

### 3.3. H3K79 methylation level is closely associated with Trpc6 expression level

We observed that in *Dot1L* KO progenitor cells, *Trpc6* level decreased, and we further investigated whether *Trpc6* expression level correlates with H3K79 di- and tri-methylation status. We chose mouse bone marrow cells as high *Trpc6* expressing and liver cells as low *Trpc6* expressing cells (**Fig. 3A**). RT-PCR and western blot analyses confirmed the expression levels of *Trpc6* in bone marrow and liver cells (**Fig. 3A** and **B**). ChIP-qPCR assays were performed for H3K79me2 and H3K79me3 by using PCR primers designed to cover the whole *Trpc6* locus, including promoter region, transcription start site, middle and end of the gene loci (**Fig. 3C**). Our results demonstrated a significant enrichment of H3K79 di- and tri-methylation at the promoter region and transcription start site of *Trpc6* in the bone marrow cells (**Fig. 3D** and **E**). In the middle and end regions of the *Trpc6* gene locus, enrichment of H3K79 methylation in bone marrow cells was not significantly higher than that of liver cells (**Fig. 3D** and **E**).

**Fig. 3.**
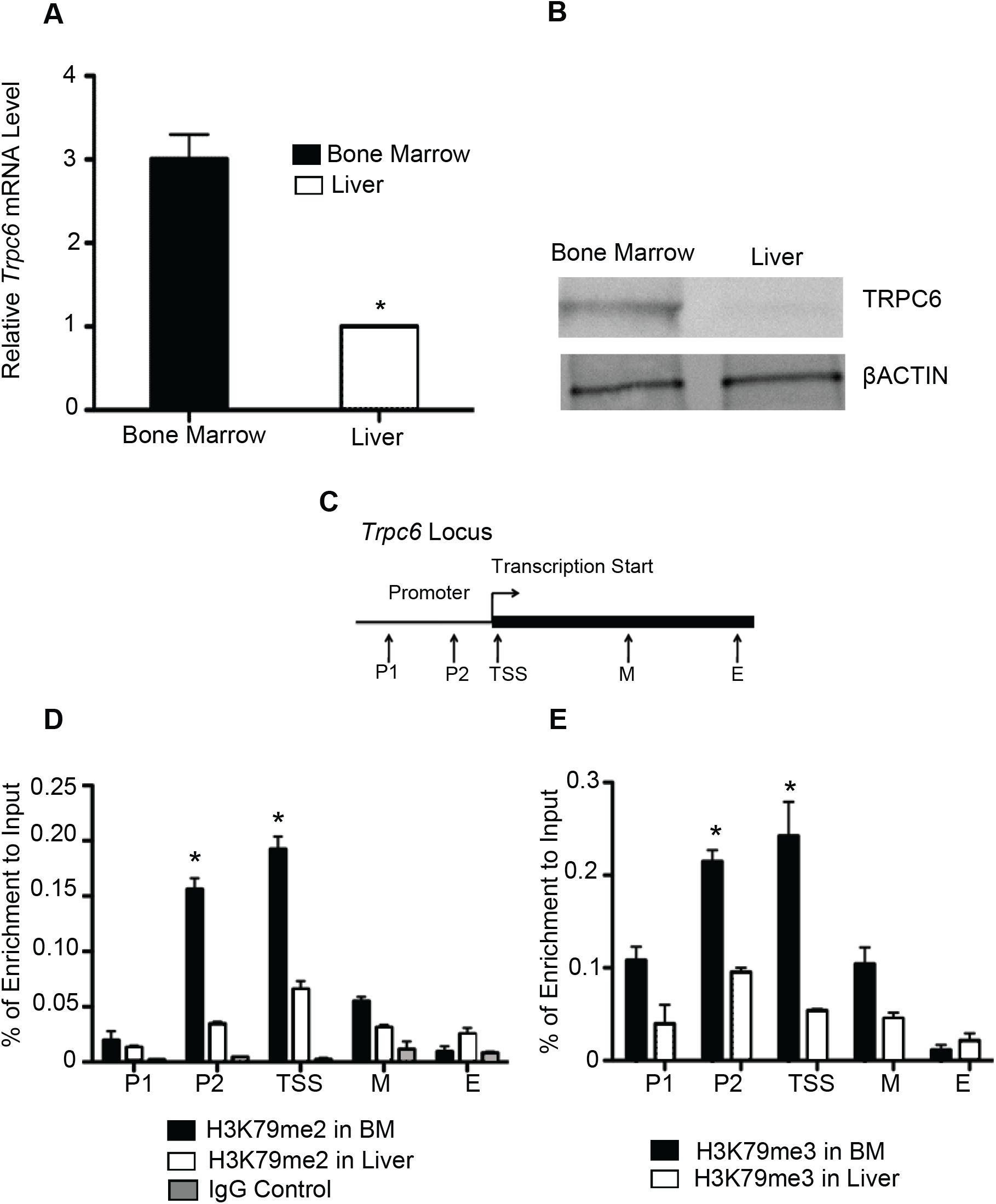
Enrichment of di- and tri-methyl H3K79 in *Trpc6* gene loci. *Trpc6* expression was tested by RT-PCR in bone marrow and liver cells. Expression of *Trpc6* was found to be significantly higher in bone marrow compared to liver (**A**). TRPC6 protein was tested by Western Blot in bone marrow and liver cells, which showed that TRPC6 was expressed in bone marrow but was barely detectable in liver cells (**B**). ChIP was performed on chromatin from bone marrow (BM) and liver cells to assess H3K79 di- and tri-methylation in *Trpc6* locus. (**C**) Several pairs of primers covering the *Trpc6* locus were designed, and H3K79 methylation in the promoter region (P1, P2), transcription start site (TSS), middle and end of gene locus (M, E) was analyzed. H3K79 dimethylation (**D**) and tri-methylation (**E**) were examined in *Trpc6* locus. There was significant enrichment of H3K79 di- and tri-methylation at the promoter region and transcription start site of *Trpc6* in bone marrow cells. The data represent three independent experiments. Data are expressed as mean±SE, n ≥ 6, * P< 0.05.

### 3.4. Calcium influx is abnormal in Dot1L-KO hematopoietic progenitor cells in response to EPO

We have tested whether Ca^2+^influx is affected in *Dot1L*-KO progenitor cells upon EPO treatment. We used the E10.5 yolk sac as the source of progenitor cells and analyzed the c-KIT positive cells for Ca^2+^ influx during a period of 20 minutes using a fluorescence microscopy-coupled digital imaging system (**Fig. 4**). We observed that when the wildtype yolk sac cells are exposed to EPO, the intracellular Ca^2+^ concentration starts to increase gradually after 5 minutes and reaches a plateau approximately after 15 minutes. In sharp contrast, the Ca^2+^ levels of *Dot1L*-KO yolk sac cells increased immediately after EPO treatment and continued increasing throughout the observation period. After 20 minutes, the Ca^2+^ signal reached a level twice as strong as that of the wildtype yolk sac cells (**Fig. 4**).

**Fig. 4.**
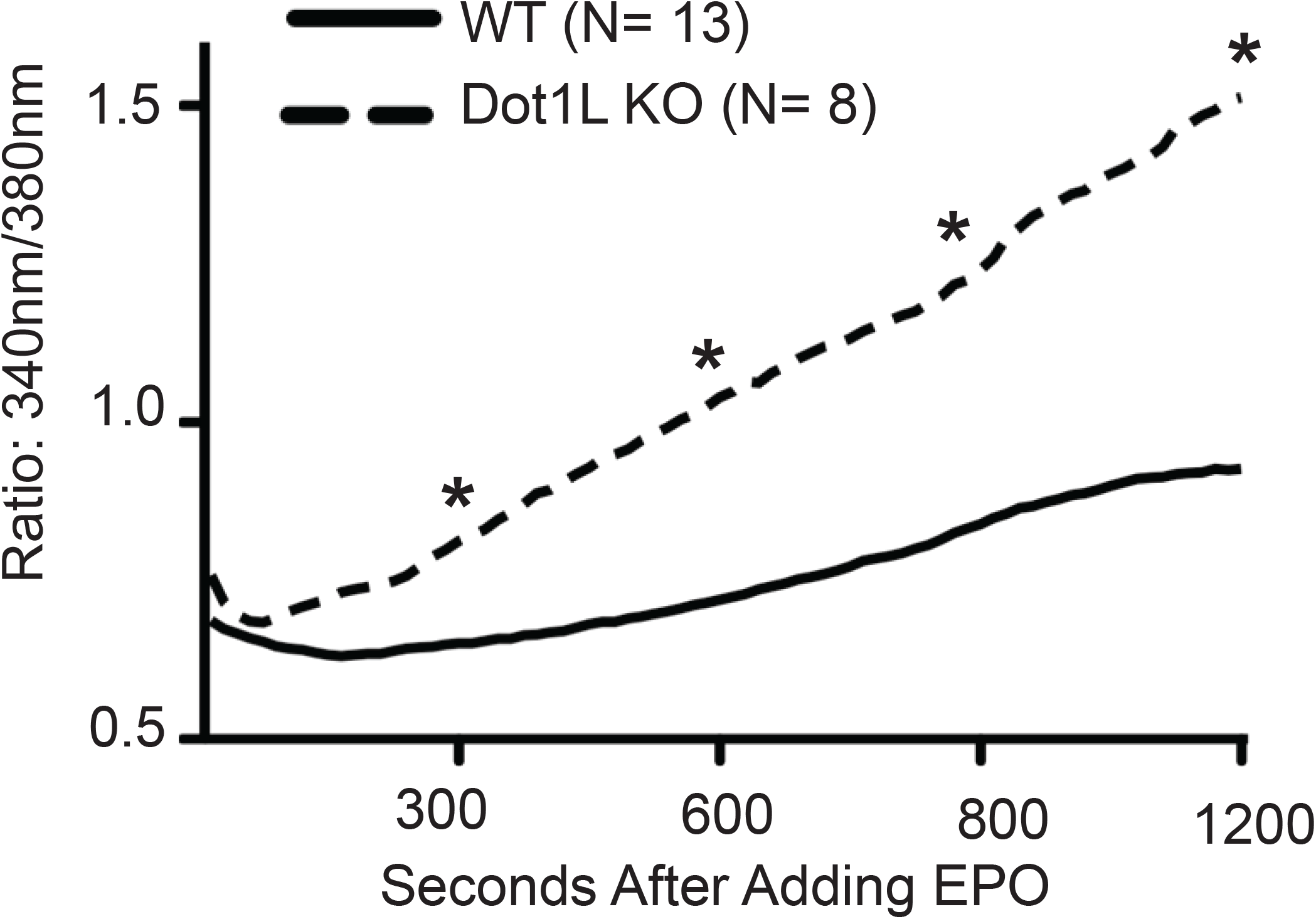
Accelerated and enhanced calcium influx in Dot1L-KO erythroid progenitors. E10.5 *Dot1L* KO and WT yolk sac cells were stained, treated with EPO, and calcium signal in individual cell was recorded by fluorescent microscopy. After EPO treatment, progenitor cells from KO yolk sacs showed sustained increase in calcium levels. Data are expressed as mean±SE, n ≥ 6, * P< 0.05.

## 4. DISCUSSION

We have observed that the first wave of primitive hematopoietic stem cells as well as the second wave of erythroid progenitor cells are present in the yolk sacs despite the loss of DOT1L protein in mutant embryos. Our findings also showed that the lack of DOT1L gives rise to a condition where these progenitors fail to generate required number of erythroid cells to support the survival of the growing fetus. We observed that an *ex vivo* culture of HPCs from wildtype yolk sacs with EPO based cytokine combinations results in a huge proliferation of erythroid progenitor cells. In contrast, the progenitor cells from *Dot1L*-KO yolk sacs undergo cell cycle arrest and apoptosis in the same culture conditions. These findings suggest that stimulation with EPO leads to death of *Dot1L*-KO erythroid progenitor cells instead of self-renewal. This study identified dysregulation of a crucial gene, *Trpc6*, in *Dot1L*-KO erythroid progenitor cells, which negatively regulates EPO-dependent Ca^2+^influx.

Both mRNA and protein levels of TRPC6 were reduced in c-KIT positive *Dot1L*-KO hematopoietic progenitor cells. We observed that expression of *Trpc6* correlates with the level of *Dot1L* expression (or H3K79 methylation). We also detected an enrichment of di- and tri-methyl H3K79 in the promoter and proximal regions of the *Trpc6* gene loci near the transcription start site. DOT1L is the only known methyltransferase that meditates H3K79 methylation in mammalian cells. Based on these findings, we can conclude that *Trpc6* is direct target of DOT1L in erythroid progenitor cells.

TRPC proteins play an important role in regulating the rate and state of Ca^2+^ influx (Gees et al., 2010). Among the *Trpc* family members, only the expression of *Trpc6* was downregulated in *Dot1L-*KO erythroid progenitor cells, while others including *Trpc2* levels remained intact, resulting in aa four-fold increase in TRPC2/TRPC6 ratio. Previous studies have shown that interaction between TRPC2 and TRPC6 is essential for maintaining EPO induced Ca^2+^ signaling (Chu et al., 2004). Consequently, we observed an accelerated and increased level of sustained Ca^2+^entry in *Dot1L-*KO erythroid progenitor cells.

Previous studies have shown that EPO signaling results in Ca^2+^ influx, which plays an important role in cell survival, proliferation and regulation of differentiation. However, dysregulation of intracellular Ca^2+^ levels due to disruption of TRPC6 expression may result in toxic effects. EPO induced calcium influx induces protein kinase activation that mediates hematopoiesis. However, excessive high levels of calcium influx can result in sustained levels of PI3K, ERK, and AKT activation, which may lead to cell cycle arrest or apoptosis. Thus, a reduced level of TRPC6 expression due to loss of DOT1L in erythroid progenitor cells may result in lethal anemia despite normal EPO signaling.

## ACKNOWLEDGEMENTS

This work was supported by the National Institutes of Health (grant R01DK091277). The mouse model was generated in the Transgenic and Gene Targeting Institutional Facility of the University of Kansas Medical Center, supported in part by the Center of Biomedical Research Excellence (COBRE) Program Project in Molecular Regulation of Cell Development and Differentiation (NIH P30 GM122731) and the University of Kansas Cancer Center (NIH P30 CA168524).

